# Ecological plant epigenetics: Evidence from model and non-model species, and the way forward

**DOI:** 10.1101/130708

**Authors:** Christina L. Richards, Conchita Alonso, Claude Becker, Oliver Bossdorf, Etienne Bucher, Maria Colomé-Tatché, Walter Durka, Jan Engelhardt, Bence Gaspar, Andreas Gogol-Döring, Ivo Grosse, Thomas P. van Gurp, Katrin Heer, Ilkka Kronholm, Christian Lampei, Vít Latzel, Marie Mirouze, Lars Opgenoorth, Ovidiu Paun, Sonja Prohaska, Stefan A. Rensing, Peter Stadler, Emiliano Trucchi, Kristian Ullrich, Koen J. F. Verhoeven

## Abstract

Growing evidence makes a strong case that epigenetic mechanisms contribute to complex traits, with implications across many fields of biology from dissecting developmental processes to understanding aspects of human health and disease. In ecology, recent studies have merged ecological experimental design with epigenetic analyses to elucidate the contribution of epigenetics to plant phenotypes, stress response, adaptation to habitat, or species range distributions. While there has been some progress in revealing the role of epigenetics in ecological processes, many studies with non-model species have so far been limited to describing broad patterns based on anonymous markers of DNA methylation. In contrast, studies with model species have benefited from powerful genomic resources, which allow for a more mechanistic understanding but have limited ecological realism. To understand the true significance of epigenetics for plant ecology and evolution, we must combine both approaches transferring knowledge and methods from model-species research to genomes of evolutionarily divergent species, and examining responses to complex natural environments at a more mechanistic level. This requires transforming genomics tools specifically for studying non-model species, which is challenging given the large and often polyploid genomes of plants. Collaboration between molecular epigeneticists, ecologists and bioinformaticians promises to enhance our understanding of the mutual links between genome function and ecological processes.

## Introduction

The idea that epigenetic variation may be important for the ecology and evolution of species has captivated biologists during recent years. The term “epigenetics” has been defined in several ways, here we focus on DNA methylation which is the mechanism that has been most studied. Earlier evidence of natural variation in DNA methylation, as well as of the inheritance and phenotypic effects of this epigenetic variation (e.g. Cubas *et al*. 1999), led to several conceptual papers that stressed its potential relevance to ecology and evolution (e.g. Richards 2006; Bossdorf *et al*. 2008; Jablonka & Raz 2009; Richards *et al*. 2010), and empirical work has been catching up slowly. Ecologists and evolutionary biologists are particularly interested in the unique contributions that epigenetic mechanisms might make. First, environment-sensitive epigenetic mechanisms could transmit responses to environmental triggers across generation boundaries. Second, heritable epigenetic variants that arise stochastically can affect phenotypes that may be under selection. In principle, this might contribute an epigenetic component to adaptation, or it might affect the dynamics of genetically based adaptation, independently from DNA sequence variation.

Research in ecological and evolutionary epigenetics is concerned with (A) patterns of natural epigenetic variation, (B) the origins and drivers of this variation, and (C) its ecological and evolutionary consequences (Bossdorf *et al*. 2008). Understanding these patterns, causes and consequences requires insight into a number of key questions that span research fields from molecular biology to ecology (Fig. 1): (A) What is the extent and structure of epigenetic variation in natural populations? (B1) What is the interplay between genetic variation and epigenetic variation? (B2) How frequently do spontaneous epimutations occur and how stable are they? (B3) To what extent can environmental changes induce heritable epigenetic changes? (C1) What is the relative importance of genetic versus epigenetic variation in determining phenotypes? (C2) How important is epigenetic variation for biotic interactions, biodiversity, and the structure and functioning of communities and ecosystems? (C3) Does epigenetic variation play a role in adaptation, and the evolution of populations?

**Figure 1.**
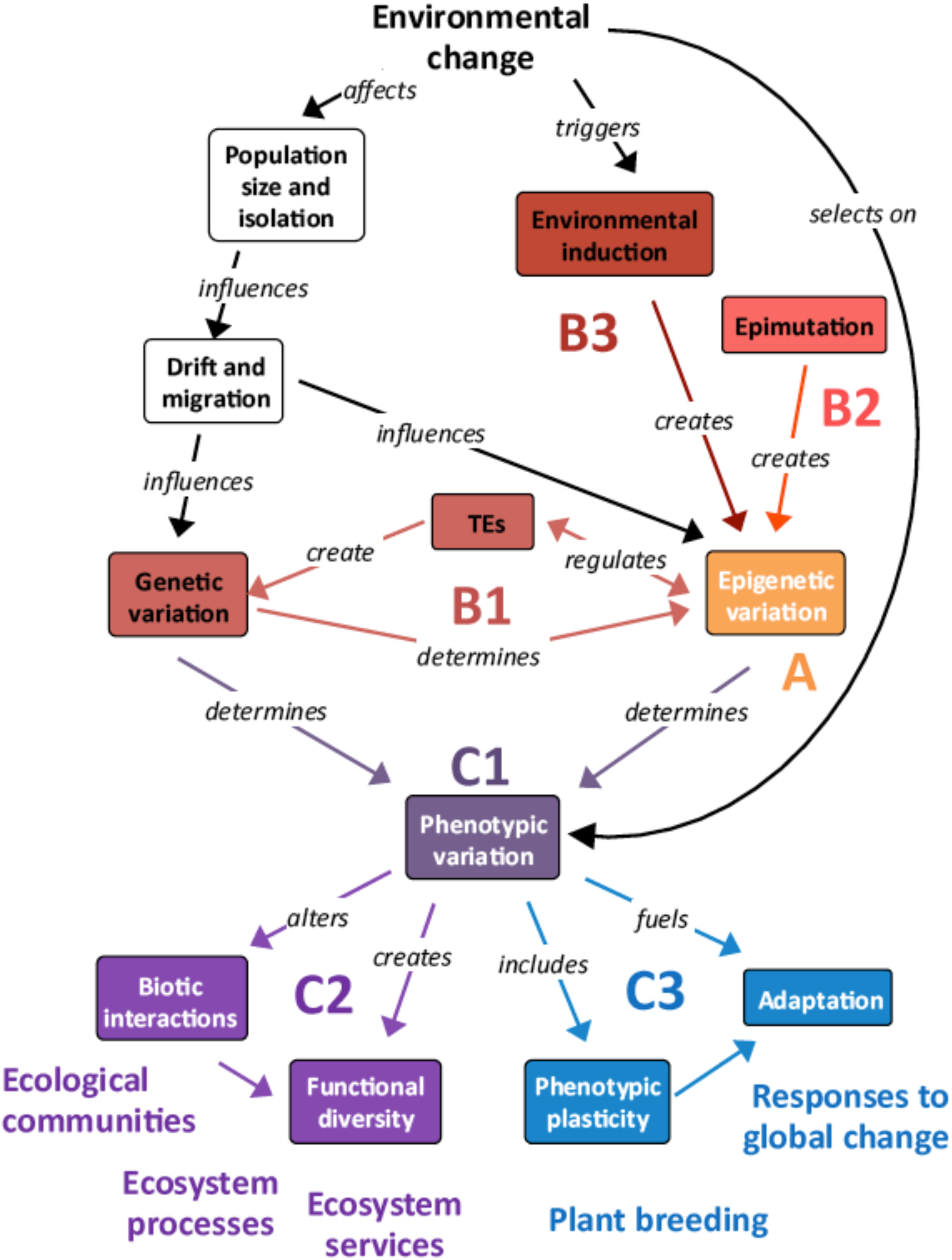
Research in ecological and evolutionary epigenetics is concerned with (A) patterns of natural epigenetic variation, (B) the origins and drivers of this variation, and (C) its phenotypic, ecological, and evolutionary consequences. Text outside of colored boxes indicate additional contributions and down stream effects of these sources of variation that contribute to the seven key questions outlined in the text. Here, “Environmental change” includes habitat reduction and fragmentation, which are primary causes of declines in population size, and can result in selection on phenotypic variation within populations.

Thus far, these questions have been studied rather separately in two different research fields: molecular genetics and evolutionary ecology. Molecular genetics has made progress in understanding the underlying mechanisms and dynamics of epigenetic variation by applying high-resolution genomic analysis tools to model species like *Arabidopsis thaliana, Oryza sativa* and *Zea mays*. Evolutionary ecology on the other hand has started to explore epigenetic variation in a broad range of non-model species that lack extensive genomic resources, and in ecological settings. This adds biological realism and has uncovered correlations between epigenetic markers with ecologically relevant variation. However, lack of genomic resources has limited the use of high-resolution tools in most non-model plant species making it difficult to firmly establish the mechanistic links between genotype, epigenotype, phenotype and environment.

To advance the study of ecological and evolutionary epigenetics, we should better integrate the fields of molecular genetics and evolutionary ecology, by adding ecological realism and ecological questions to model species research (e.g. Latzel *et al*. 2013; Hagmann *et al*. 2015), and by adopting higher-resolution tools in non-model species research (e.g. Platt *et al*. 2015; Xie *et al*. 2015; Grugger *et al*. 2016; van Gurp *et al*. 2016; Trucchi *et al*. 2016). A similar transition took place in ecological and evolutionary genomics (Pavey *et al*. 2012; Narum *et al*. 2013; Alvarez *et al*. 2015), which revealed the importance of ecological realism in study designs because organismal responses under laboratory conditions may not reflect performance under natural conditions, and that non-adaptive processes may result in apparent genomic signatures of selection (Pavey *et al*. 2012; Alvarez *et al*. 2015).

This review focuses on progress accomplished by molecular epigenetics and evolutionary ecology in plants, following the series of seven key questions outlined above. DNA methylation is currently the most frequently studied and best-understood epigenetic process, so our review is largely restricted to studies of DNA methylation. However, it is important to note that the relevance of histone modifications and small RNAs that are involved in guiding epigenetic modifications is increasingly acknowledged (see e.g., Kim *et al*. 2015; Matzke & Mosher 2014), and the potential for interconnection among different epigenetic mechanisms is not yet fully understood (Becker *et al*. 2011). Here, we seek to identify next steps, discussing strategies and methodological challenges in future ecological epigenetics research.

### Current progress in ecological epigenetics-What have we learned?

#### A. What are the patterns of natural epigenetic variation?

What is the extent and structure of epigenetic variation in natural populations?

Global cytosine methylation varies widely within and among plant species (4– 40%; Fig. 2; Alonso *et al*. 2014, 2015; Niederhuth *et al*. 2016). Full genome bisulfite sequencing has only been performed in model plants, and studies in *A. thaliana* (Schmitz *et al*. 2013; Kawakatsu *et al*. 2016) and *O. sativa* (He *et al*. 2010; Chodavarapu *et al*. 2012; Li *et al*. 2012) found variable methylation among accessions, which depends on sequence context (i.e. CG, CHG, or CHH), and genomic regions (gene promoters, gene bodies, transposable elements (TEs)). While detailed information on genomic context is lacking for non-model species, studies in wild plant populations have documented extensive intraspecific variation in DNA methylation, based on anonymous markers (Schrey *et al*. 2013), global DNA methylation estimates (Alonso *et al*. 2015), and assessment at specific genes (Xie *et al*. 2015; see Box 1).

**Figure 2.**
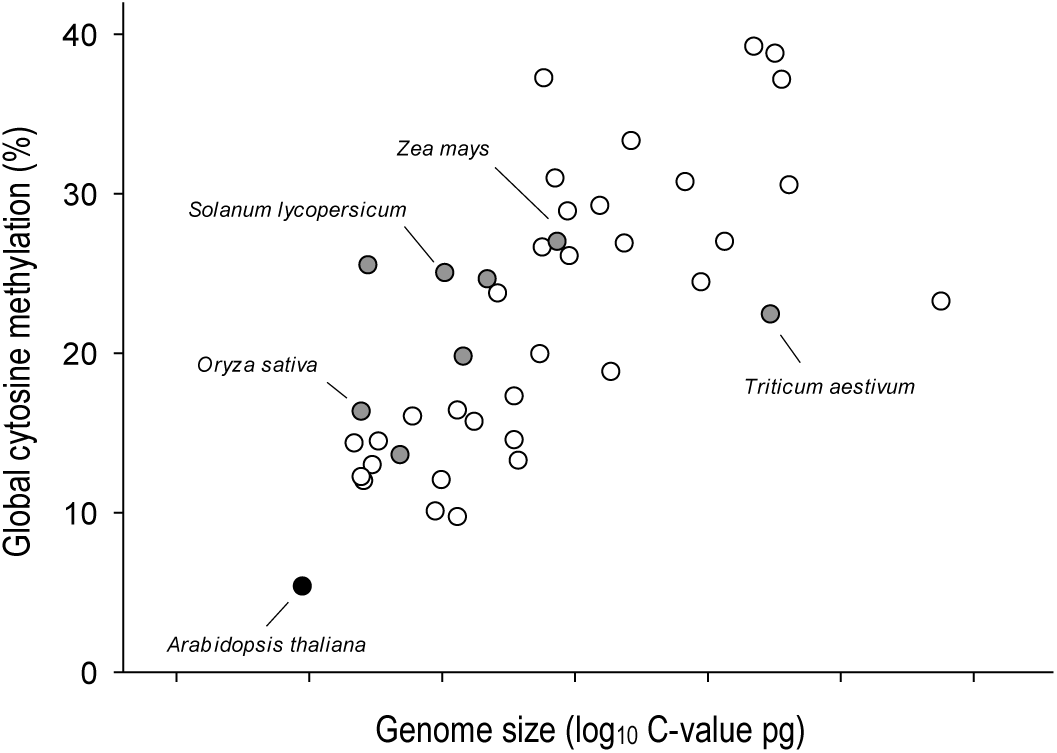
Features and limitations of *Arabidopsis thaliana* as a model system for epigenetic studies. Rapid development, selfing ability, and reduced chromosome number and genome size (C-value = 0.30 pg) with relatively few repetitive and transposable elements, facilitate experimentation and simplify molecular analyses in *A. thaliana*. However, *A. thaliana* (black dot) is unusual within the range of variation in genome size and global cytosine methylation in study species, including some with a fully sequenced reference genome (filled dots) like rice (*Oryza sativa*), tomato (*Solanum lycopersicum*), maize (*Zea mays*) and wheat (*Triticum aestivum*). Redrawn from Alonso *et al*. (2015).

###### 1. Global methylation

Global levels of DNA methylation can be estimated through HPLC-and ELISA-based assays that provide an estimate of the proportion of methylated cytosines. These methods do not require any sequence knowledge, and they do not distinguish the genomic location or genetic context (CG, CHG, CHH) of 5mC. Nevertheless, they can help to clarify the magnitude of variation in global methylation across species (Alonso *et al*. 2015), the structuring of natural intraspecific variation (Alonso *et al*. 2014), or the global response to specific treatments.

###### 2. Methylation-sensitive markers

Genome-wide patterns of DNA methylation can be captured by molecular markers obtained with methylation-sensitive restriction enzymes. **Methylation-sensitive AFLP** (Reyna-Lopez *et al*. 1997; MS-AFLP / MSAP) follows a standard AFLP protocol but uses isoschizomers with differential sensitivity to DNA methylation, often *HpaII* and *MspI*. MS-AFLP typically evaluates several hundreds of restriction sites. AFLP and MS-AFLP can be applied in parallel to examine how the environment structures genetic and epigenetic variation (Herrera & Bazaga 2010; Richards *et al*. 2012; Schulz *et al*. 2014; Foust *et al*. 2016; Robertson *et al*. in press). MS-AFLP has been a popular approach in ecological epigenetics in previous years largely due to easy application to non-model organisms and the fact that no reference genome or advanced bioinformatics skills are required. However, the method is gradually being replaced by bisulfite sequencing-based methods. A new method, **EpiRADseq** (Schield *et al*. 2015), combines methylation-sensitive restriction enzymes with NGS. Similar to MS-AFLP, the method detects methylation differences only in the recognition sequence, but it produces a much greater number of loci and sequence information at interrogated loci.

###### 3. Bisulfite sequencing methods

Bisulfite treatment converts unmethylated cytosines to uracil, providing the basis for identifying methylated cytosines upon sequencing and comparing a treated sample to a proper reference or untreated sample (Cokus *et al*. 2008). This is the gold standard of DNA methylation analysis, which can evaluate each cytosine in the target sequence of candidate loci or can provide quantitative methylation information for essentially all cytosines in a genome (i.e. **whole-genome bisulfite sequencing** or **WGBS**). While WGBS enables detailed functional and genetic analysis of DNA methylation variation and its environmental or transgenerational dynamics, it is restricted to species with a high-quality reference genome, and its costs may be prohibitive for large sample sizes and in species with large genomes. This currently limits its use for ecological epigenetic studies involving non-model species, and ecological experimental designs. Bisulfite sequencing can be applied more broadly when restricting sequencing to specific subsets of the genome. If a high-quality reference transcriptome is available, bisulfite sequencing combined with **exome capture** permits the interrogation of methylation status of the expressed regions of the genome (and their flanking regions; Lee *et al*. 2011).

Likewise, bisulfite sequencing can be targeted to a selection of genomic fragments created with restriction enzymes (**Reduced representation bisulfite sequencing**, **RRBS**; Gu *et al*. 2011), providing nucleotide resolution of DNA methylation within each of the isolated fragments. The availability of both sequence and methylation variation from the same large set of loci (for instance, thousands to tens of thousands at 10x coverage; Narum *et al*. 2013) provides for the direct comparison of genetic and epigenetic information, allowing evaluation of the relative contribution of SNPs and DMPs to population divergence. Moreover, availability of sequence context for identified methylation polymorphisms opens the door to the functional analysis required for causally linking DNA methylation variation to phenotypic variation, especially in the context of population divergence and local adaptation. RRBS has been adopted for plant population studies (Platt *et al*. 2015), and methods have been developed to incorporate bisulfite sequencing into popular reduced representation sequencing approaches that can be applied also to species for which no reference genome is available (bsRADseq and epiGBS; Trucchi *et al*. 2016; van Gurp *et al*. 2016).

###### 4. ChIP-Sequencing

Chromatin immunoprecipitation followed by next-generation sequencing (ChIP-seq) determines the modification state of histone proteins (Park *et al*. 2009). Specific antibodies bind to the histone modification of interest and immonuprecipitate fragments of DNA that are wrapped around the modified histones. These DNA fragments are sequenced and mapped back to the reference genome to determine the specific regions where the modification was present.

###### 5. sRNA-sequencing

Comparative data on the diversity and abundance of small RNAs across individuals can be obtained by strand-specific deep sequencing of small RNA molecules (Davey *et al*. 2010). Whereas differential abundance across environments or treatments can readily be tested after clustering the sRNA reads per individual, functional interpretations are aided by a reference genome or transcriptome.

Given that DNA methylation variation can result from genetic control, environmental induction and stochastic epimutations, and can in principle be shaped further by drift and selection, interpretation of patterns of natural epigenetic variation is not straightforward. Ecological studies have searched for epigenetic variation that correlates with habitats or with population differentiation, and report that variation in DNA methylation usually exceeds variation in genetic markers when populations from contrasting habitats are compared (e.g., Lira-Medeiros *et al*. 2010; Herrera & Bazaga 2010; Richards *et al*. 2012; Medrano *et al*. 2014; Schulz *et al*. 2014; but see Foust *et al*. 2016; Robertson *et al. in press*). Further, DNA methylation often correlates with ecological factors in a way that is not predicted from the observed underlying patterns of genetic relatedness (Richards *et al*. 2012; Schulz *et al*. 2014; Foust *et al*. 2016, Gugger *et al*. 2016). However, how much of the correlation between DNA methylation and different environmental factors *in situ* is repeatedly induced versus stably inherited, or under direct genetic control, is still an open question.

#### B. What are the origins and drivers of epigenetic variation?

##### B1 What is the interplay between genetic variation and epigenetic variation?

Epigenetic variants may contribute to heritable trait variation, but if the epigenetic variants are entirely under genetic control then the implications of epigenetic variation reduces to the underlying genetics (Richards 2006). Genetic control of epigenetic variation is indicated by work in *A. thaliana* that has revealed how sequence changes in genes related to the epigenetic machinery (e.g. methyltransferases *MEE57* and *CMT2*; Becker *et al*. 2011; Dubin *et al*. 2015) can have dramatic effects on the epigenome (see below under “What can be transferred from model to non-model species?“). Quantitative genetic studies in model plant species also suggest that many DNA methylation differences between individuals are associated with underlying genetic variants. For example, the majority of differentially methylated regions (DMRs) stably segregated with the local allele in maize (*cis* inheritance; Li *et al*. 2014). In contrast, only 35% of DMRs were associated with genetic polymorphisms among 142 natural *A. thaliana* accessions (Schmitz *et al*. 2013). Associations between DMRs and genetic polymorphisms in *cis* or in *trans* can indicate genetic control over DNA methylation. However, *cis* associations can also arise in the absence of genetic control, when a spontaneous epimutation is stably inherited through epigenetic inheritance (Taudt *et al*. 2016).

On the other hand, epigenetic change can also feed back on genetic variation since DNA methylation is associated with silencing of transposable elements. Reduction of DNA methylation can result in the up-regulation of TE expression and transposition, potentially creating new genetic variants and phenotypes when integrating near endogenous genes, attracting repressive epigenetic marks and influencing gene expression. In fact, many of the well-characterized epialleles are associated with TEs or repeats that can cause DNA methylation in *cis* or in *trans* via RNA-directed DNA methylation (RdDM; Paszkowski & Grossniklaus 2011; O’Malley & Ecker 2012; Matzke & Mosher 2014).

Studies in non-model plants have attempted to isolate the effects of epigenetic variation while controlling for genetic diversity by using species that have naturally low levels of genotypic diversity such as asexually reproducing plants (e.g. Verhoeven *et al*. 2010; Richards *et al*. 2012; Verhoeven & Preite 2014). In outcrossing species, a few studies using molecular markers have used statistical approaches to make inferences about the independence of genetic and epigenetic variation by evaluating how well overall similarities in DNA methylation profiles between individuals can be predicted from their genetic similarity (Herrera and Bazaga 2010; Schulz *et al*. 2014; Foust *et al*. 2016). Although these approaches in non-model species have provided circumstantial evidence for epigenetic variation that may be independent from genetic control, these studies cannot rule out the possibility that genetic polymorphisms have gone unnoticed and generated epigenetic variants (Becker *et al*. 2011; Dubin *et al*. 2015).

##### B2. How often do spontaneous epimutations occur?

Stochastic epimutations have the potential to contribute to heritable trait variation in a way that is not predictable from underlying sequence variation, but their evolutionary potential is determined by the rate at which they appear and revert. Epimutation rates have been assessed in *A. thaliana* mutation accumulation lines, where DNA methylation polymorphisms accumulated at a much higher frequency (van der Graaf *et al*. 2015) than genetic mutations calculated for the same population (Ossowski *et al*. 2010). These differentially methylated positions (DMPs) occurred more frequently in genic regions than in TEs consistent with the known RdDM mechanism, which limits development of methylation polymorphisms in these contexts (Teixeira *et al*. 2009). Spontaneous epimutations also occurred at the level of DMRs, which showed more functional relevance to gene expression, but also occurred at much lower frequency, comparable to the rate at which SNPs arise (Becker *et al*. 2011).

Data on epimutation rates are absent for non-model plant species. However, in apomictic dandelions, Verhoeven *et al*. (2010) showed that even in a constant environment, appreciable DNA methylation differences developed between individual plants, and that most of these changes were inherited to apomictically produced offspring. Further studies are needed to determine the rate and stability of newly arising epialleles and DMRs, and to understand their functional consequence on gene expression, traits, and fitness, and therefore their potential role in adaptation.

##### B3. To what extent can environmental changes induce heritable epigenetic changes?

Studies in *A. thaliana* have shown that the epigenome reacts to external influences such as abiotic and biotic stress (e.g. Dowen *et al*. 2012; Slaughter *et al*. 2012; Sani *et al*. 2013), and reports have shown that these epigenetic changes can sometimes be associated with changes in gene expression throughout the genome (Secco *et al*. 2015; Wibowo *et al*. 2016). Further, some studies have reported stress-induced transgenerational DNA methylation effects (e.g. in rice: Kou *et al*. 2011; in *A. thaliana*: Bilichak *et al*. 2012). In particular, herbivore or pathogen effects can persist into offspring generations, and some of the best current evidence for epigenetic-mediated transgenerational effects is in the context of such biotic interactions in *A. thaliana* (Luna *et al*. 2012; Rasmann *et al*. 2012). However, another study showed that methylation changes in response to hyperosmotic stress were inherited for only one generation, and methylation changes were largely reset in the absence of the stress (Wibowo *et al*. 2016). Other reports in *A. thaliana* and maize have detected very limited stress-induced DNA methylation modifications that were transmitted to offspring (Pecinka & Mittelsten Scheid 2012; Eichten & Springer 2015).

Studies in non-model species have found that heritable changes in methylation status of specific markers occur in response to different environmental stresses, but the responsiveness of methylation depends on environmental conditions (Verhoeven *et al*. 2010; Alonso *et al*. 2016). For instance, Verhoeven *et al*. (2010) showed that plants with identical genotypes exposed to different stresses had a range of changes in methylation-sensitive markers in response to different stresses, and the rate of transmission of these changes to offspring also varied. Richards *et al*. (2012) measured epigenetic differences in a dominant haplotype of the highly invasive clonal plant Japanese knotweed (*Fallopia spp.*) collected from different habitats, but grown in a common environment. They found that methylation of different DNA markers correlated with habitat of origin, suggesting environment-induced methylation changes that persisted through clonal propagation. Similarly, after growing the invasive plant *Ageratina adenophora* under controlled conditions, Xie *et al*. (2015) found that stable inheritance of demethylation at the promoter region of a specific gene was correlated to geographic variation in cold tolerance. However, whether these studies reflect stable, environmentally induced methylation patterns or selection of variants with different methylation profiles remains to be determined.

#### C. What are the consequences of epigenetic variation?

##### C1. What is the contribution of DNA methylation to phenotypes?

So far, only a few natural epialleles have been functionally characterized using various approaches (Box 2; Cubas *et al*. 1999; Manning *et al*. 2006; Paszkowski & Grossniklaus 2011; Xie *et al*. 2015). Model plants offer powerful tools to isolate epigenetic effects from genetic effects on phenotype. Of particular note are two populations of epigenetic recombinant inbred lines (epiRILs) from *A. thaliana*, derived from crosses between the Columbia wild type and mutants in the Columbia background that have decreased DNA methylation genome wide (*ddm1* and *met1* mutants; Johannes *et al*. 2009; Reinders *et al*. 2009; see below in “What can be transferred from model to non-model species?“). Both *met1* and *ddm1* epiRIL populations displayed significant phenotypic variation (Zhang *et al*. 2013; Cortijo *et al*. 2014), and linkage mapping in the *ddm1* epiRILs identified segregating DMRs that explained heritable phenotypic effects for root length and flowering time (QTL^epi^, Cortijo *et al*. 2014).

###### Box 2: Approaches to link epigenetic variation to phenotype

Establishing the link between epigenetic variation and phenotype is one of the key questions in ecological epigenetics.

###### Candidate genes

Detailed observational studies can reveal epigenetic polymorphisms at candidate loci that are tightly associated with expression and phenotypic differences in a genetically uniform background (such as completely inbred lines, asexually propagated individuals, or within the same individual, e.g. Cubas *et al*. 1999; Xie *et al*. 2015). Such observations also suggest autonomous epigenetic determinants of phenotypic variation.

###### Epigenetic mutants

Mutants that are compromised in an epigenetic mechanism, such as de novo or maintenance of DNA methylation, can cause genome-wide alterations in epigenetic marks, which can be exploited to investigate whether phenotypes are influenced by epigenetic mechanisms. Mutants also have been useful in experiments where the environmental induction and transmission of epigenetic changes is investigated. If a certain environment cannot induce phenotypic changes in mutant background (Rasmun *et al*. 2012; Luna *et al*. 2012) or transmission of a phenotype is altered (Crevillen *et al*. 2014) this allows for identifying the epigenetic mechanism responsible.

###### Chemical manipulation

Mutants are not often available in non-model organisms, but certain drugs can be used to manipulate the epigenetic variation such as 5-azacytidine (Jones 1985) or zebularine (Cheng *et al*. 2003) that act as DNA methylation inhibitors. There are also chemicals that target different epigenetic mechanisms, such as inhibition of H3K27me3 demethylation (Kruidenier *et al*. 2012) However, these chemicals can have potentially cytotoxic or other off-target effects, and the chemicals can be biased to specific loci (Hagemann *et al*. 2011).

In ecological studies, DNA methylation inhibitors have been used to show variation among genotypes in phenotypic responses to different environments, and the importance of DNA methylation for plant responses to environmental conditions (Bossdorf *et al*. 2010, Herrera *et al*. 2012), maintaining the effects of inbreeding (Vergeer *et al*. 2012), flowering time differences (Wilschut *et al*. 2016), and inheritance of induced phenotypes (Herman *et al*. 2016).

###### Epigenetic association and QTL-mapping

Ecologists are most interested in natural variation, and in principle the same methods of quantitative genetics that are applicable for the study of natural genetic variation can be used to investigate natural epigenetic variation. Several studies have screened for genetic markers that contribute to differences in methylation patterns by treating methylation as a phenotype (e.g. Dubin *et al*. 2015), which helps to unravel genetically controlled versus autonomous DNA methylation variation. Conversely, DMPs or DMRs that are stably inherited across generations can be treated the same way as conventional genetic markers to permit mapping approaches. Long *et al*. (2011) used both genetic and methylation markers for QTL-mapping in a *Brassica napus* population, as many of the methylation states segregated in a normal Mendelian fashion, which was also observed for DMRs in a maize pedigree (Li *et al*. 2014). Epigenetic markers have been used for QTL analysis in plants in the absence of DNA sequence polymorphisms (Cortijo *et al*. 2014). We are not aware of any studies that use such markers for association mapping in non-model species other than in some low-resolution methylation sensitive AFLP studies with plants. These marker-based mapping studies can shed light on the epigenetic contribution to phenotypic variation. However, a statistical epigenetic marker-trait association does not necessarily imply a true epigenetic contribution to the observed trait variation, as a phenotype may be caused by a tightly linked genetic polymorphism. Follow-up studies are required to functionally characterize the suspected QTL^epi^ and to evaluate it in further manipulative studies.

In non-model species, several studies have found correlations between anonymous methylation-sensitive AFLP markers and leaf traits (Herrera & Bazaga 2013), flower morphology (Herrera & Bazaga 2010) and fitness-related traits (Medrano *et al*. 2014) in natural populations. While it is tempting to interpret these correlations in terms of contributions to adaptation, they may be simply a read-out of genetic differences. Identification of the responsible loci will be required for confirmation of the causal relationships between epigenetic variation and phenotypes.

##### C2. What are the ecological consequences of epigenetic variation?

By mediating plastic responses, DNA methylation can facilitate response to various biotic and abiotic conditions, and persistence in different environments. For instance, using in vivo demethylation by the methylation inhibitor azacytidine, Herrera *et al*. (2012) showed that the exploitation of nectar of varying sugar concentrations in a flower-living yeast depended on DNA methylation. Similarly, Herrera and Bazaga (2011) found both DNA sequence and methylation polymorphisms in *Viola cazorlensis* were correlated with herbivory damage and habitat. In addition, variation in genomic DNA methylation has been correlated to shifts in species range (Richards *et al*. 2012; Xie *et al*. 2015), and differentiation of populations (Platt *et al*. 2015; Foust *et al*. 2016; but see Robertson *et al*. *in press*). Interactions between plants and biotic or abiotic stressors can prime plants for a more rapid or vigorous response upon a second exposure in the future (Conrath *et al*. 2002), and epigenetic mechanisms such as histone modifications may be responsible for such sustained stress memory (e.g. Jaskiewicz *et al*. 2011).

In the model plant *A. thaliana*, there is some evidence suggesting that the epigenetic contribution to response to ecologically important factors like nutrient availability depended on genotype among natural accessions (Bossdorf *et al*. 2010). Experiments with epiRILs demonstrated that epigenetic variation among lines created functional diversity that had very similar effects on population and ecosystem functioning as found for genetic and species diversity effects: higher epigenetic variation created variation in phenotypes that translated into increased productivity and resistance of experimental populations (Latzel *et al*. 2013). The combination of studies in natural accessions and epiRILs suggests that epigenetic diversity may be an important component of functional biodiversity, and that epigenetic variation can be indirectly involved in evolution by modifying selection at the community level.

##### C3. What are the evolutionary consequences of epigenetic variation?

Several studies have investigated the role of environmentally induced and spontaneous epigenetic modifications in evolutionary theory (Jablonka & Raz 2009; Slatkin 2009; Day & Bonduriansky 2011; Geoghegan & Spencer 2012; Klironomos *et al*. 2013; Charlesworth & Jain 2014; Furrow 2014; Wang & Fan 2014; Kronholm & Collins 2016). Ultimately, the impact of spontaneous epigenetic variation in evolution will depend on the rates and stability of epigenetic changes and the distribution of their phenotypic effects (Kronholm & Collins 2016). Modelling studies show that if spontaneous epigenetic changes occur at faster rates than genetic changes, this could lead to evolutionary dynamics where phenotypic changes are first driven by epigenetic changes, and become genetically encoded later (Klironomos *et al*. 2013; Kronholm & Collins 2016). These studies have also shown that environmentally induced epigenetic changes that are inherited across generations should be adaptive in rapidly changing environments (Robertson & Richards 2015). However, the potential significance of such environmentally induced variation for adaptation will strongly depend on its persistence and stability. Detailed analyses in *A. thaliana*, have revealed that seemingly adaptive patterns of natural epigenetic variation may be under genetic control (Dubin *et al*. 2015), and that little evidence exists for environment-induced epigenetic variation that persists for several generations (Hagmann *et al*. 2015).

### What can be transferred from model to non-model species?

The transfer of information from model systems combined with advances in sequencing and bioinformatics approaches will offer a powerful next step for ecological epigenetics due to more precise insights into function and an increase in genome coverage. Some of the detailed information on epigenetic mechanisms that we have learned from model species is already useful in non-model systems. Gene annotations in model species provide information on genes that code for conserved components of the epigenetic machinery or genes that are epigenetically regulated. Studies of plant RdDM (Khraiwesh *et al*. 2010), CMT methyltransferase (Noy-Malka *et al*. 2014) and other epigenetic marks like histone H3K27 trimethylation demonstrate that epigenetic mechanisms are controlled by evolutionarily conserved machinery (Rensing *et al*. 2008; Widiez *et al*. 2014). This information can be used to identify homologs in non-model species, which can then be targets for knock-outs or transformation to validate function in future studies (Kobayashi *et al*. 2013; Alvarez *et al*. 2015; Xie *et al*. 2015).

Studies in *A. thaliana* have shown that mutations occurring in genes related to the epigenetic machinery can have strong effects on epigenetic variation. In a study of near-isogenic mutation accumulation lines, a single spontaneous mutation in *MEE57*, a MET1-related methyltransferase, presumably led to an increased rate of epimutation at CG sites, resulting in 40% more methylation differences after 30 generations compared to the other MA lines (Becker *et al*. 2011). In another recent study on natural populations, Dubin *et al*. (2015) found a link between alternative alleles of the DNA methyltransferase *CMT2* (responsible for CHG and CHH methylation of certain classes of TEs) and the epigenome’s capacity to respond to temperature changes. In contrast, the CMT3 homolog double mutants in maize are not viable (Li *et al*. 2014), indicating that loss of methylation can have more drastic effects in some species.

Further studies in *A. thaliana* have shown that the vast majority of small RNAs (more than 60%) are of the 24 nt size class (Kasschau *et al*. 2007), and complement sequences that are homologous to TEs and are involved in the targeting of DNA methylation through RdDM (Matzke & Mosher 2014). For instance, small RNAs target DNA methylation to long terminal repeats (LTRs) of retrotransposons (where the promoter is located), and inhibit TE transcription. Reduction of DNA methylation at that location can be associated with the upregulation of TE expression, potentially creating new genetic variants and phenotypes. Studies in RdDM deficient *A. thaliana* have shown that TEs get activated and introduce new copies in the genome (Mirouze *et al*. 2009; Ito *et al*. 2011). Thus, RdDM could be an important mechanism that protects the genome from TE amplification. TEs can also play an important role when they integrate near endogenous genes, and influence gene expression. A tight interplay exists between TEs, the anti-TE activity of RdDM and epigenetic regulation of gene expression.

In addition to the detailed information about genes involved in the epigenetic machinery, two research groups created epigenetic recombinant inbred lines (epiRILs) to specifically isolate epigenetic from genetic information (question B1), and to be able to study the dynamics and phenotypic consequences of DNA methylation (C1) in the almost complete absence of DNA sequence variation. The epiRILs were created from crosses between *A. thaliana* mutants with decreased genome-wide levels of DNA methylation (*ddm1* and *met1*; Johannes *et al*. 2009; Reinders *et al*. 2009) and their wild-type counterparts. The progeny was either backcrossed or selfed, and only individuals homozygous for the wild type DDM1 and MET1 allele were allowed to self-fertilize for multiple generations to increase epi-homozygosity of segregating epialleles. The populations of epiRILs are therefore nearly isogenic but segregate hundreds of stably hypomethylated regions from the mutant parent, allowing for an assessment of the phenotypic consequences of epigenetic variation independent of variation in DNA sequence. Studies with the epiRIL populations have shed light on the mechanisms involved in inheritance of DNA methylation and the phenotypic impact of epigenomic alterations. First, DNA methylation is in some genomic regions inherited in a stable Mendelian fashion (Colomé-Tatché *et al*. 2012). In the *ddm1* epiRIL population, linkage mapping identified segregating DMRs that explained observed heritable phenotypic effects (Cortijo *et al*. 2014). This demonstrated, for the first time in any organism, that DMRs can be stably inherited for many generations independently of DNA sequence and that they can act as epigenetic quantitative trait loci (QTL^epi^). From a phenotypic evolution point of view these QTL^epi^ have all the necessary properties to become targets of natural or artificial selection.

Further studies with *ddm1* epiRILs demonstrated that some DMRs showed patterns of non-Mendelian inheritance, mainly through the action of small RNAs especially at TE loci (Teixeira *et al*. 2009). In addition, the *ddm1* epiRILs showed heritable variation in phenotypic plasticity and stress tolerance (Zhang *et al*. 2013), and that epigenetically diverse plant populations were more productive and more stable than epigenetically uniform populations (Latzel *et al*. 2013). In the *met1* epiRIL population, some aberrant phenotypes were linked to TE mobilization (Mirouze *et al*. 2009): certain hypomethylated TEs segregated in the population and proliferated until they were eventually silenced again through post-transcriptional and then transcriptional mechanisms (Marí-Ordóñez *et al*. 2013). The *ddm1* mutation also caused transposition of previously silenced retrotransposons (Miura *et al*. 2001).

Despite all of the insight from studies of model plants, there are limitations to the transfer of knowledge to non-model species that we must keep in mind when we try to adapt these findings to non-model species. In particular, the genome and epigenome of *A. thaliana* are atypical for most plants that have been surveyed (Fig. 2; Alonso *et al*. 2015). Studies across diverse taxa have demonstrated that TEs can be tightly linked to epigenetic regulation of plant gene expression, but the comparatively small *A. thaliana* genome has relatively few TEs, which are mainly clustered around the centromeres. Larger plant genomes contain proportionally more TEs and they are typically more evenly distributed. For example, the 500 Mbp *Physcomitrella patens* moss genome contains 50% TEs with no discernible peak regions (Rensing *et al*. 2008). Recent analysis has indicated that even among plants with small genomes, *A. thaliana* might constitute an “epigenetic outlier” due to its conspicuously reduced number of TEs: the genome of the close relative *A. lyrata* has a considerably larger TE density and consequently contains more methylated sites and regions (Seymour *et al*. 2014). In addition, TEs are not well conserved evolutionarily even between closely related species (Seymour *et al*. 2014), so insights on causes of methylation changes identified in one species may not always transfer to another. Further, plant genomes differ even in the classes of TEs that they harbor. For example, nucleocytoplasmic large DNA virus (NCLDV) insertions have only been shown in non-seed plant genomes (Maumus *et al*. 2014), and the maize genome is subject to reshuffling specifically by Helitrons (rolling-circle replicating transposons), that are estimated to be involved in expression of 25% of the maize transcriptome (Barbaglia *et al*. 2012). In studies of other epigenetic mechanisms, the repressing histone H3K9 methylation varies with genome size. Whereas small genomes such as *A. thaliana* show H3K9 methylation in constitutive heterochromatin, genomes of 500 Mbp and larger display H3K9 methylation in euchromatic regions as well (Houben *et al*. 2003). Analyses of epigenetic mechanisms in species with bigger and more complex genomes, including other model and non-model species, will allow for generality and contribute to better understanding the relevance of epigenetic mechanisms for plant adaptation and evolution.

### Future directions

Ultimately, ecological epigenetics studies are interested in understanding the epigenetic contributions to ecological and evolutionary processes in nature. High-resolution genomic analyses particularly in *A. thaliana* have provided concrete evidence for epigenetic variants with heritable phenotypic effects, the selective potential of empirically observed epimutation rates, and the inheritance of environment-induced epigenetic effects. However, some studies suggest that natural DNA methylation variation in *A. thaliana* is largely under genetic control and that there is little evidence of epigenetic legacies from environments experienced in previous generations. In contrast, in non-model species, a variety of studies show that DNA methylation variation correlates with ecological factors in a way that is not simply predicted from underlying genetic relatedness, suggesting a genetics-independent role for epigenetics. But, the low genomic resolution of many studies precludes pinpointing causality of epigenetic effects.

A priority for ecological epigenetics in the coming years is therefore to incorporate tools that enable higher-resolution analysis in more realistic scenarios, and in a wider diversity of systems. This is important because the role of epigenetics is expected to vary between species, with different genomic features and ploidy levels (Ainouche *et al*. 2009; Niederhuth *et al*. 2016; Springer *et al*. 2016; Takuno *et al*. 2016). Furthermore, selection may have shaped the genetic capacity for transgenerational epigenetic effects differently in different environments and species because the selective advantages of phenotypic plasticity and transgenerational effects differ between species depending on habitat predictability and life history characteristics (Herman *et al*. 2014; Verhoeven & Preite 2014).

Addressing the questions related to epigenetic contributions to ecological and evolutionary processes will continue to be challenging not only because of the tools required to tie functionality to epigenetic changes, but also the complicated relationship between epigenetic variation and DNA sequence variation, and the labile nature of epigenetic variation (Richards *et al*. 2010). Pinpointing causality of autonomous epigenetic variation is challenging even in model systems, and studies that extend analysis into ecological settings and non-model species face a trade-off between precision, realism and generality of results (Levins 1966). With better tools available, what kind of evidence should we expect from future ecological epigenetics studies?

### Functional relevance

Field studies of epigenetics provide insight about which habitat conditions may lead to interesting patterns of epigenetic divergence, but attempts to link methylation markers to phenotype or habitat in the field cannot isolate the functional importance of epigenetics. Validation of the phenotypic effect of field-based observations of genome-wide methylation or methylation of candidate genes can be accomplished by using mutants, knockouts, or genetically engineered organisms that are altered in genes involved in the epigenetic machinery, or that are differentially methylated (e.g. epiRILs). Subsequently, the ecological relevance of the variant can be determined in classic ecological experiments in the greenhouse or field. Lacking these types of genetic resources, the broad functional importance of methylation in various ecological and evolutionary processes can be explored with chemical reduction of DNA methylation variation, which has been useful in exploring the role of DNA methylation in a variety of ecological and evolutionary processes (Box 2).

### Unraveling epigenetics and genetics

Although several authors have suggested that functional DNA methylation is largely under genetic control (e.g. Li *et al*. 2012; Dubin *et al*. 2015), we have very little data in any system that can address to what extent there is a component of epigenetic variation independent of genetic variation that contributes to organismal function. For model species, it is possible to test associations between genetic alleles and epi-alleles from QTL/EWAS mapping (e.g. Dubin *et al*. 2015), which are indicative of genetic control over epigenetic variation especially in the case of *trans* associations. In non-model species, we have described the effective use of clonal or asexual plant species for isolating the role of epigenetics, but low-level genetic variation in natural clonal lineages cannot be excluded. Considering that ecological studies are often focused on collections from natural populations *in situ*, future analyses will need to accommodate a simultaneous comparison of genetic and epigenetic data sets to examine how much of the overall epigenetic variation can be predicted from pairwise genetic relatedness, and identify differences in genetic and epigenetic patterns (Herrera & Bazaga 2010; Grugger *et al*. 2015; Lea *et al*. 2015; Foust *et al*. 2016; Herrera *et al*. 2016). These approaches have become standard using anonymous molecular markers, but approaches based on next generation sequencing will be more powerful to identify epigenetic associations with phenotype or habitat that are not predicted by the observed genetic variation. Once these associations are identified, more detailed analysis with further targeted bisulphite sequencing and/or expression of the candidate loci across different genetic backgrounds, knockouts or transgenic organisms can be used to investigate the independence and importance of the observed epigenetic effects.

### Environmental effects

Several studies have found correlations between environmental variation and epigenetic differences, leaving open the question of whether the epigenetic variation is induced or selected by the environment, or simply a side effect of genetic structuring. Environmentally-induced epigenetic effects may be either transient or persistent across generations, and heritable changes may be either selected on or linked to something that is selected. Future analyses should therefore consider that part of the epigenetic variation is similar to phenotypic variation, and carefully designed experiments are necessary to characterize both genetic and environmental contributions to epigenetic variation (Richards *et al*. 2010). Observational field surveys of natural populations can provide information on how epigenetic variation is structured on the landscape. When they simultaneously measure genetic and epigenetic data on the same individuals, statistical approaches can identify epigenetic variation that is not explained by genetic variation (Dubin *et al*. 2015; Foust *et al*. 2016). This so-called “genetic-independent” epigenetic variation may be induced transiently by the environment or it may be stably inherited. To discriminate between these two possibilities, one must grow individuals in a common garden and analyze which epigenetic marks persist. Moreover, to isolate how much epigenetic variation is induced by environment requires experiments where both genetics and environment are controlled. Here, clonal organisms or inbred lines are particularly useful because they allow for growing genetically identical offspring in different environments. In outcrossing species, different breeding designs, such as half-sibs or full sibs, and quantitative genetics mapping approaches such as EWAS can approximate genetic and epigenetic associations with phenotype.

To date, many studies of the environmental effects on epigenetic variation have tested for the existence and transgenerational persistence of environmentally induced effects, but there is limited insight into the adaptive significance of these effects. Building on theory of phenotypic plasticity (Herman *et al*. 2014), there are good *a priori* hypotheses of how the capacity for epigenetic plasticity and transgenerational effects will have different adaptive benefits depending on species life history traits and habitat characteristics, such as spatial or temporal heterogeneity and dispersal mode. Genotype-specificity in epigenetic or transgenerational effects may be common (e.g. Herman & Sultan 2016), and future ecological epigenetics studies should explore the evidence for adaptive variation in the capacity for such effects.

### Technical challenges and opportunities

While computational analyses of genome-wide data have become routine in model species, not all approaches are easily transferred to ecological epigenetics (Box 3). The main challenge for developing sequencing approaches in ecological epigenetics is that existing workflows usually require at least high quality transcriptomes, if not complete reference genomes. Two important methodological developments have the potential to advance studies in ecological epigenetics: First, recently developed approaches based on reduced-representation methods (RRBS, Box 1) allow a base-pair resolution for DNA methylation detection, and can be applied to species for which no genomic resources are available (Trucchi *et al*. 2016; van Gurp *et al*. 2016). Only short fragments of a small fraction of the genome are reconstructed in RRBS, representing only a subset of DNA methylation polymorphisms. Nevertheless, RRBS methods have successfully detected DNA methylation variation in several species (Gugger *et al*. 2016; van Gurp *et al*. 2016; Trucchi *et al*. 2016). The use of these methods in ecologically real systems will help to identify DNA methylation variants that impact performance, and motivate detailed follow-up experiments to characterize candidate loci, although there may always be undetected *trans*-acting genetic variation that controls observed DNA methylation patterns. Second, even crude draft transcriptomes and genomes support many of the standard workflows in epigenomics data analysis, therefore the additional efforts of constructing and annotating these resources will be instrumental for a more sophisticated understanding of epigenomics in non-model systems. Currently, the number of draft genomes of ecological study species is limited, but continued reduction in sequencing cost, and advances in long-read sequencing methods are rapidly bringing draft assemblies of ecological species within reach.

##### Box 3: Key bioinformatics challenges for epigenetic sequence analysis

The analysis of NGS data is a complex problem best thought of as many individual specific tasks combined in a modular fashion to form analysis pipelines or workflows. Although pipelines are still typically written as specific software, it has become easier to combine the individual components in a customized fashion using generic bioinformatic workflow management systems such as Taverna, KNIME or Galaxy.

###### Reference Construction

is a particular issue in ecological genetics and epigenetics since standard NGS analyses require complete genomic reference sequences. For work in non-model species, where no reference genome exists, researchers must use reduced-representation methods. Specialized tools have been developed that cluster nearly identical reads and then assemble them locally into reference sequences for the assayed loci (Narum *et al*. 2013; van Gurp *et al*. 2016).

###### Read Mapping

determines the source of an observed NGS sequence (the read) in a reference sequence. Many software tools are available for this task. Segemehl (Hoffmann *et al*. 2014) is optimized for large deviations between query reads and targets and can therefore be used for bisulfite-modified data. Other methods are specially designed to analyze bisulfite sequencing data, such as Bismark (Krueger & Andrews 2011) or BWA-meth (http://arxiv.org/abs/1401.1129).

###### Variation calling

identifies positions in the reference where the aligned reads deviate from the reference sequence. Variation calling includes the detection of sequence variation like SNPs, indels, and structural variants. Variation calling is integrated into software tools for mapping and reference construction and several tools for SNP calling are available (e.g. GATK, McKenna *et al*. 2010; ANGSD, Korneliussen *et al*. 2014). Some tools like Bis-SNP (Liu *et al*. 2012) and MethylExtract (Barturen *et al*. 2013) combine calling for SNPs and methylation differences from bisulfite-modified data.

###### Differential methylation calling

is the identification of differentially methylated cytosines from bisulfite converted sequencing data. Several methods have been developed to determine the differential methylation status of cytosines (Bock 2012) and are integrated into specific software (e.g. Methpipe, Bismark; Song *et al*. 2013; Krueger & Andrews 2011). However, comparative methylation analysis in model species has shown that detection of differential methylation is highly coverage-and sequence context-dependent. Since methylation rates at CHG and CHH sites are generally low, differences at these sites between samples are reliably detectable only at high sequencing coverage (>100x; Becker *et al*. 2011). In *A. thaliana*, this results in a biased detection of DMPs in CG context and gene bodies. This bias is even more pronounced in non-model species with large genome size (lower coverage) and with less reliable information on non-genic sequences.

###### Definition and identification of DMRs

Changes of DNA methylation not at individual cytosines (DMPs) but across larger chromosomal stretches (differentially methylated regions, DMRs) might be more biologically relevant, as they are more likely to influence transcriptional activity at the nearby locus and thus to contribute to phenotypic expression. Therefore, a distinction between single sites and contiguous stretches of methylation could be critical. While it has been common practice to call DMRs by either looking for clusters of DMPs (Becker *et al*. 2011) or by applying a sliding window approach over the length of the genome (Schmitz *et al*. 2013), these approaches lead to high rates of false negative and false positive calls, respectively. While DMP clusters suffer from the same bias of sequence context as single DMPs, sliding-window approaches lead to the detection of DMRs at many weakly methylated regions of the genome and the high number of false positives worsens with increasing genome size, making it not useful for many non-model species. A first generation of region callers for BS-seq data are MOABs, BSmooth, DMRcate, and metilene. A recent adaptation to plant methylation data of a Hidden-Markov-Model-based approach provides a more neutral method for DMR detection (Molero *et al*. 2011; Hagmann *et al*. 2015; Lea *et al*. 2015) by first identifying regions of dense methylation that are then tested for differential methylation; it has been successfully applied to *A. thaliana* (Hagmann *et al*. 2015), *A. lyrata* and *Capsella rubella* (Seymour *et al*. 2014), *Arabis alpina* (Willing *et al*. 2015), *P. patens*, rice and maize (Becker et al. unpublished data), proving its applicability to genomes of different sizes, and with diverse methylation distributions. While DMR calling is challenging even using WGBS data, its application to RRBS-based data from non-model species without a reference genome can be further complicated by the small size of the fragments.

###### Peak Calling

is necessary for ChIP-seq analysis and other related experimental protocols that enrich DNA or RNA from particular genomic regions. Here, statistically significant over-representation of mapped reads in particular genomic regions are identified. These local “peaks” then indicate the position of nucleosomes carrying specific chemical modifications (Bailey *et al*. 2013).

Along with the development of genomics resources, there is a need also for further development of data analysis methods. For instance, identifying proper statistical testing approaches for differential DNA methylation in complex ecological experimental designs (including random effects like population and genotype) is an ongoing challenge even for DMPs (e.g. Lea *et al*. 2015). The definition and identification of DMRs holds additional challenges, particularly for RRBS fragments (which are often only the length of a so-called “region“) and for DNA methylation in different genomic contexts. Existing software tools do not yet cope with these complexities (Box 3). Further, while DNA methylation has been the most studied epigenetic modification in the context of natural variation, histone methylation marks also can be inherited (Gaydos *et al*. 2014; Ragunathan *et al*. 2014; Audergon *et al*. 2015), and there is evidence for small RNA-dependent epialleles (Calo *et al*. 2014). Thus, epigenetic mechanisms other than DNA methylation need to be investigated in an ecological context, too.

Since organisms in natural settings are continuously exposed to multiple environmental signals and must respond appropriately to dynamic conditions, an ecological context provides a unique opportunity to discover information about epigenetic variation that cannot be gleaned through controlled laboratory settings. Recent studies in natural settings have found gene expression patterns that are only exposed under complex natural stimuli (Pavey *et al*. 2012; Alvarez *et al*. 2015), but the role of epigenetics is almost unexplored. Studies in non-model organisms may also yield functional information about genomic elements that are not annotated in model species, have no homolog in their most closely-related model organism, or have taken on a novel function. Applying new tools and understanding of epigenetics and genome function in general to a robust ecological design will be powerful for assessing the importance of both genetic and epigenetic mechanisms in the real world.

## Acknowledgements

This work arose from the workshop ‘sEpiDiv - Towards understanding the causes and consequences of epigenetic diversity’ organized by K.H. and L.O. The workshop was held at and kindly supported by the Synthesis Centre of the German Centre for Integrative Biodiversity Research (iDiv) Halle-Jena-Leipzig (DFG FZT 118). All authors contributed to the discussions and participated in drafting the manuscript, and the writing was partially supported by funding from the National Science Foundation DEB-1419960 (to CLR).

## References

Ainouche, M. L., Fortune, P. M., Salmon, A., Parisod, C., Grandbastien, M. A., Fukunaga, K., Ricou, M. & Misset, M-T. (2009). Hybridization, polyploidy and invasion: lessons from Spartina (Poaceae). Biological invasions 11.5, 1159–1173.

Alonso, C., Pérez, R., Bazaga, P. & Herrera, C. M. (2015). Global DNA cytosine methylation as an evolving trait: phylogenetic signal and correlated evolution with genome size in angiosperms. Front. in Genet. 6, 4.

Alonso, C., Pérez, R., Bazaga, P., Medrano, M. & Herrera, C. M. (2014). Individual variation in size and fecundity is correlated with differences in global DNA cytosine methylation in the perennial herb Helleborus foetidus (Ranunculaceae). Am. J. Bot. 101, 1309–1313.

Alonso, C., Pérez, R., Bazaga, P., Medrano, M. & Herrera, C. M. (2016). MSAP markers and global cytosine methylation in plants: a literature survey and comparative analysis for a wild growing species. Mol. Ecol. Res. 16, 80–90 doi: 10.1111/1755-0998.12426.

Alvarez, M., Schrey, A. W. & Richards, C. L. (2015). Ten years of transcriptomics in wild populations: what have we learned about their ecology and evolution? Mol. Ecol. 24, 710–725.

Audergon, P. N. C. B., Catania, S., Kagansky, A., Tong, P., Shukla, M., Pidoux, A. L. & Allshire, R. C. (2015). Epigenetics. Restricted epigenetic inheritance of H3K9 methylation. Science 348, 132–135.

Bailey, P. K., Ladunga, I., Lefebvre, C., Li, Q., Liu, T., Madrigal, P., Taslim, C. & Zhang, J. (2013). Practical guidelines for the comprehensive analysis of ChIP-seq data. PLoS Comp. Biol. 9(11), e1003326. doi: 10.1371/journal.pcbi.100332.

Barbaglia, A. M., Klusman, K. M., Higgins, J., Shaw, J. R., Hannah, L. C. & Lal, S. K. (2012). Gene capture by Helitron transposons reshuffles the transcriptome of maize. Genetics 190, 965–975.

Barturen, G., Rueda, A., Oliver, J. L. & Hackenberg, M. (2014). MethylExtract: High-Quality methylation maps and SNV calling from whole genome bisulfite sequencing data. Version 2. F1000Res. 2013 Oct 15 [revised 2014 Feb 21], 2, 217. doi: 10.12688/f1000research.2-217.v2. eCollection.

Becker, C., Hagmann, J., Müller, J., Koenig, D., Stegle, O., Borgwardt, K. & Weigel, D. (2011). Spontaneous epigenetic variation in the Arabidopsis thaliana methylome. Nature 480, 245–249.

Bilichak, A., Ilnystkyy, Y., Hollunder, J. & Kovalchuk, I. (2012). The progeny of *Arabidopsis thaliana* plants exposed to salt exhibit changes in DNA Methylation, histone modifications and gene expression. PLoS ONE 7(1): e30515.

Bock, C. (2012). Analysing and interpreting DNA methylation data. Nature Rev. Genet. 13, 705–719.

Bossdorf, O., Arcurri, D., Richards, C. L. & Pigliucci, M. (2010). Experimental alteration of DNA methylation affects the phenotypic plasticity of ecologically relevant traits in *Arabidopsis thaliana*. Evol. Ecol. 24, 541–553.

Bossdorf, O., Richards, C. L. & Pigliucci, M. (2008). Epigenetics for ecologists. Ecol. Lett. 11, 106–115 (2008).

Calo, S., Shertz-Wall, C., Lee, S. C., Bastidas, R. J., Nicolas, F. E., Granek, J. A., Mieczkowski, P., Torres-Martinez, S., Ruiz-Vazquez, R. M., Cardenas, M. E. & Heitman, J. (2014). Antifungal drug resistance evoked via RNAi-dependent epimutations. Nature 513, 555–558.

Charlesworth, B. & Jain, K. (2014). Purifying selection, drift, and reversible mutation with arbitrarily high mutation rates. Genetics 198(4), 1587–602.

Cheng, J. C., Matsen, C. B., Gonzales, F. A., Ye, W., Greer, S., Marquez, V. E., Jones, P. A. & Selker, E. U. (2003). Inhibition of DNA methylation and reactivation of silenced genes by Zebularine. Journal of the National Cancer Institute 95, 399–409.

Chodavarapu, R. K., Feng, S., Ding, B., Simon, S. A., Lopez, D., Jia, Y., Wang, G. L., Meyers, B. C., Jacobsen, S. E. & Pellegrini, M. (2012). Transcriptome and methylome interactions in rice hybrids. Proc. Natl Acad. Sci. USA 109, 12040– 12045.

Cokus, S. J., Feng, S., Zhang, X., Chen, Z., Merriman, B., et al. (2008). Shotgun bisulphite sequencing of the *Arabidopsis* genome reveals DNA methylation patterning. Nature 452, 215–219.

Colomé-Tatché, M., Cortijo, S., Wardenaar, R., Lahouze, B., Etcheverry, M., Martin, A., Feng, S., Duvernois-Berthet, E., Labadie, K., Wincker, P., Jacobsen, S. E., Jansen, R. C., Colot, V. & Johannes, F. (2012). Features of the *Arabidopsis* recombination landscape resulting from the combined loss of sequence variation and DNA methylation. Proc. Natl. Acad. Sci. USA 149, 16249–16245.

Conrath, U., Pieterse, C. M. J. & Mauch-Mani, B. (2002). Priming in plant–pathogen interactions. Trends in Plant Sci. 7 (5), 210–216.

Cortijo, S., Wardenaar, R., Colomé-Tatché, M., Gilly, A., Etcheverry, M., Labadie, K., Caillieux, E., Hospital, F., Aury, J.-M., Wincker, P., Roudier, F., Jansen, R. C., Colot, V. & Johannes, F. (2014). Mapping the epigenetic basis of complex traits. Science 343(6175), 1145–1148.

Crevillen, P., Yang, H., Cui, X., Greeff, C., Trick, M., Qiu, Q., Cao, X. & Dean, C. (2014). Epigenetic reprogramming that prevents transgenerational inheritance of the vernalized state. Nature 515, 587–590.

Cubas, P., Vincent, C. & Coen, E. (1999). An epigenetic mutation responsible for natural variation in floral symmetry. Nature 401, 157–161.

Davey, J. W. & Blaxter, M. L. (2010). RADSeq: next-generation population genetics. Brief. Funct. Genomics 9, 416–423.

Day T. & Bonduriansky, R. (2011). A unified approach to evolutionary consequences of genetic and nongenetic inheritance. Am. Nat. 178, E18–E36.

Dowen, R. H., Pelizzola, M., Schmitz, R. J., Lister, R., Dowen, J. M., Nery, J. R., Dixon, J. E. & Ecker, J. R. (2012). Widespread dynamic DNA methylation in response to biotic stress. Proc. Natl. Acad. Sci. USA 109, E2183–2191.

Dubin, M. J., Zhang, P., Meng, D., Remigereau, M.-S., Osborne, E. J., Casale, F. P., Drewe, P., Kahles, A., Vilhjálmsson, B., Jagoda, J., Irez, S., Voronin, V., Song, Q., Long, Q., Rätsch, G., Stegle, O., Clark, R. M. & Nordborg, M. (2015). DNA methylation variation in Arabidopsis has a genetic basis and shows evidence of local adaptation. eLife 4.

Eichten, S. R. & Springer, N. M. (2015). Minimal evidence for consistent changes in maize DNA methylation patterns following environmental stress. Front. Plant Sci. 6, 308.

Foust, C. M., Preite, V., Schrey, A. W., Verhoeven, K. J. F. & Richards, C. L. (2016). Genetic and epigenetic differences associated with environmental gradients in replicate populations of two salt marsh perennials. Mol. Ecol. 25, 1639–1652.

Furrow, R.E. (2014). Epigenetic inheritance, epimutation, and the response to selection. PLoS One 9(7): e101559.

Gaydos, L. J., Wang, W. & Strome, S. (2014). H3K27me and PRC2 transmit a memory of repression across generations and during development. Science 345, 1515–1518.

Geoghegan, J.L., Spencer, H.G. (2012). Population-epigenetic models of selection. Theor. Popul. Biol. 81(3), 232–42.

van der Graaf, A., Wardenaar, R., Neumann, D. A., Taudt, A., Shaw, R. G., Jansen, R. C., Schmitz, R. J., Colomé-Tatché, M. & Johannes, F. (2015). Rate, spectrum and evolutionary dynamics of spontaneous epimutations. Proc. Natl. Acad. Sci. USA 112, 6676–6681.

Gu, H., Smith, Z. D., Bock, C., Boyle, P., Gnirke, A. & Meissner, A. (2011). Preparation of reduced representation bisulfite sequencing libraries for genome-scale DNA methylation profiling. Nature Protoc. 6(4), 468–481.

Gugger, P., Fitz-Gibbon, S., Pellegrini, M. & Sork, V.L. (2016). Species wide patterns of DNA methylation variation in Quercus lobata and its association with climate gradients. Mol. Ecol. 25, 1665–1680.

van Gurp, T. P., Wagemaker, N. C. A. M., Wouters, B., Vergeer, P., Ouborg, J. N. J. & Verhoeven, K. J. F. (2016). epiGBS: reference-free reduced representation bisulfite sequencing. Nat. Meth. 13, 322–324.

Hagemann, S., Heil, O., Lyko, F. & Brueckner, B. (2011). Azacytidine and decitabine induce gene-specific and non-random DNA demethylation in human cancer cell lines. PLoS ONE 6, e17388.

Hagmann, J., Becker, C., Müller, J., Stegle, O., Meyer, R. C., Wang, G., Schneeberger, K., Fitz, J., Altmann, T., Bergelson, J., Borgwardt, K. & Weigel, D. (2015). Century-scale methylome stability in a recently diverged Arabidopsis thaliana lineage. PLoS Genet. 11(1), e1004920.

He, G., Zhu, X., Elling, A. A., Chen, L., Wang, X., Guo, L., Liang, M., He, H., Zhang, H. Chen, F. et al. (2010). Global epigenetic and transcriptional trends among two rice subspecies and their reciprocal hybrids. Plant Cell 22, 17–33.

Herman J.J. & Sultan, S.E. (2016). DNA methylation mediates genetic variation for adaptive transgenerational plasticity. Proc Roy Soc B 283, doi:10.1098/rspb.2016.0988.

Herman, J. J., Spencer, H. G., Donohue, K. & Sultan, S.E. (2014). How stable ‘should’ epigenetic modifications be? Insights from adaptive plasticity and bet hedging. Evolution 68, 632–643.

Herrera, C. M. & Bazaga P. (2011). Untangling individual variation in natural populations: ecological, genetic and epigenetic correlates of long-term inequality in herbivory. Mol Ecol. 20, 1675–1688.

Herrera, C. M. & Bazaga, P. (2013). Epigenetic correlates of plant phenotypic plasticity: DNA methylation differs between prickly and nonprickly leaves in heterophyllous *Ilex aquifolium* (Aquifoliaceae) trees. Bot. J. Linean Soc. 171, 441–452.

Herrera, C. M. & Bazaga, P. (2010). Epigenetic differentiation and relationship to adaptive genetic divergence in discrete populations of the violet *Viola cazorlensis*. New Phytol. 187(3), 867–876.

Herrera, C. M., Medrano, M. & Bazaga, P. (2016). Comparative spatial genetics and epigenetics of plant populations: heuristic value and a proof of concept. Mol. Ecol. 25, 1653–1664.

Herrera, C. M., Pozo, M. I. & Bazaga, P. Jack of all nectars, master of most: DNA methylation and the epigenetic basis of niche width in a flower-living yeast. Mol. Ecol. 21, 2602–2616 (2012).

Hoffmann, S., Otto, C., Doose, G., Tanzer, A., Langenberger, D., Christ, S., Kunz, M., Holdt, L. M., Teupser, D., Hackermüller, J. & Stadler, P. F. (2014). A multi-split mapping algorithm for circular RNA, splicing, *trans*-splicing and fusion detection. Genome Biol. 15(2), R34. doi: 10.1186/gb-2014-15-2-r34.

Houben, A., Demidov, D., Gernand, D., Meister, A., Leach, C.R. & Schubert, I. (2003). Methylation of histone H3 in euchromatin of plant chromosomes depends on basic nuclear DNA content. The Plant Journal: for cell and molecular biology 33, 967–973.

Ito, H., Gaubert, H., Bucher, E., Mirouze, M., Vaillant, I. & Paszkowski, J. (2011). An siRNA pathway prevents transgenerational retrotransposition in plants subjected to stress. Nature 472, 115–119.

Jablonka, E. B. & Raz, G. (2009). Transgenerational epigenetic inheritance: prevalence, mechanisms, and implications for the study of heredity and evolution. Q. Rev. Biol. 84(2), 131–176.

Jaskiewicz, M. et al. (2011). Chromatin modification acts as a memory for systemic acquired resistance in the plant stress response. EMBO Rep. 12, 50–55.

Johannes, F., Porcher, E., Teixeira, F., Saliba-Colombani, V., Simon, M., Agier, N., Bulski, A., Albuisson, J., Heredia, F., Bouchez, D., Dillmann, C., Guerche, P., Hospital, F. & Colot, V. (2009). Assessing the impact of transgenerational epigenetic variation on complex traits. PLoS Genetics 5, e1000530.

Jones, P. A. (1985). Altering gene expression with 5-azacytidine. Cell 40, 485–486.

Kasschau, K. D., Fahlgren, N., Chapman, E. J., Sullivan, C. M., Cumbie, J. S., Givan, S. A. & Carrington, J. C. (2007). Genome-wide profiling and analysis of *Arabidopsis* siRNAs. PLoS Biol. 5, e57.

Kawakatsu, T., Huang, S. C., Jupe, F., Sasaki, E., Schmitz, R. J., Urich, M. A., Castanon, R., Nery, J. R., Barragan, C., He, Y., Chen, H., Dubin, M., Lee, C.-R., Wang, C., Bemm, F., Becker, C., O’Neil, R., O’Malley, R. C., Quarless, D. X., The 1001 Genomes Consortium, Schork, N. J., Weigel, D., Nordborg, M. & Ecker, J. R. (2016). Epigenomic diversity in a global collection of *Arabidopsis thaliana* accessions. Cell 166, 492–505.

Khraiwesh, B., Arif, M. A., Seumel, G. I., Ossowski, S., Weigel, D., Reski, R. & Frank, W. (2010). Transcriptional control of gene expression by microRNAs. Cell 140, 111–122.

Kim, J.-M., Sasaki, T., Ueda, M., Sako, K. & Seki, M. (2015). Chromatin changes in response to drought, salinity, heat, and cold stresses in plants. Front. in Pl. Sci. 6, 114.

Klironomos F., Berg, J. & Collins, S. (2013). How epigenetic mutations can affect genetic evolution: Model and mechanism. BioEssays 35, 571–578.

Kobayashi, M. J., Takeuchi, Y., Kenta, T., Kume, T., Diway, B. & Shimizu, K. K. (2013). Mass flowering of the tropical tree *Shorea beccariana* was preceded by expression changes in flowering and drought-responsive genes. Mol. Ecol. 22, 4767–4782.

Korneliussen, T. S., Anders, A. & Nielsen, R. (2014). “ANGSD: analysis of next generation sequencing data.” BMC bioinformatics 15.1: 356

Kou, H., Li, Y., Song, X., Ou, X., Xing, S., Ma, J., Von Wettstein, D. & Liu, B. (2011). Heritable alteration in DNA methylation induced by nitrogen-deficiency stress accompanies enhanced tolerance by progenies to the stress in rice (*Oryza sativa* L.) J. of Pl. Physiol. 168, 1685–1693.

Kronholm, I. & Collins, S. (2016). Epigenetic mutations can both help and hinder adaptive evolution. Mol. Ecol. 25, 1856–1868.

Krueger, F. & Andrews, S. R. (2011). Bismark: a flexible aligner and methylation caller for Bisulfite-Seq applications. Bioinformatics 27(11), 1571–1572.

Kruidenier, L., Chung, C.-W., Cheng, Z., Liddle, J., Che, K., Joberty, G., Bantscheff, M., Bountra, C., Bridges, A., Diallo, H., Eberhard, D., Hutchinson, S., Jones, E., Katso, R., Leveridge, M., Mander, P. K., Mosley, J., Ramirez-Molina, C., Rowland, P., Schofield, C. J., Sheppard, R. J., Smith, J. E., Swales, C., Tanner, R., Thomas, P., Tumber, A., Drewes, G., Oppermann, U., Patel, D. J., Lee, K. & Wilson, D. M. (2012). A selective jumonji H3K27 demethylase inhibitor modulates the proinflammatory macrophage response. Nature 488, 404–408.

Latzel, V., Allan, E., Silveira, A. B., Colot, V., Fischer, M. & Bossdorf, O. (2013). Epigenetic diversity increases the productivity and stability of plant populations. Nat. Commun. 4, 2875.

Lea, A.J., Tung, J. & Zhou, X. (2015). A flexible, efficient binomial mixed model for identifying differential DNA methylation in bisulfite sequencing data. PLoS Genet. 11(11): e1005650.

Lee, E. J., Pei, L., Srivastava, G., Joshi, T., Kushwaha, G., Choi, J. H., et al. (2011). Targeted bisulfite sequencing by solution hybrid selection and massively parallel sequencing. Nucl. Acids Res. 39(19), e127–e127.

Levins R. (1966). The strategy of model building in population biology. Am. Sci. 54, 421–431.

Li, Q., Eichten, S. R., Hermanson, P.J. & Springer, N.M. (2014). Inheritance Patterns and Stability of DNA methylation variation in maize near-isogenic lines. Genetics 196(3), 667–676.

Li, X., Zhu, J., Hu, F., Ge, S., Ye, M., Xiang, H., Zhang, G., Zheng, X., Zhang, H., Zhang, S., Li, Q., Luo, R., Yu, C., Yu, J., Sun, J., Zou, X., Cao, X., Xie, X., Wang, J. & Wang, W. (2012). Single-base resolution maps of cultivated and wild rice methylomes and regulatory roles of DNA methylation in plant gene expression. BMC Genomics 13, 300.

Lira-Medeiros, C. F. et al. (2010). Epigenetic variation in mangrove plants occurring in contrasting natural environment. PLoS One 5, e10326.

Liu, Y., Siegmund, K. D., Laird, P. W. & Berman, B. P. (2012) Bis-SNP: combined DNA methylation and SNP calling for Bisulfite-seq data. Genome Biology 13(7):R61

Long, Y., Xia, W., Li, R., Wang, J., Shao, M., Feng, J., King, G. J. & Meng, J. (2011). Epigenetic QTL mapping in *Brassica napus*. Genetics 189, 1093–1102.

Luna, E., Bruce, T., Roberts, M., Flors, V. & Ton, J. (2012). Next generation systemic acquired resistance. Plant Physiol. 158, 844–853.

Manning, K., Tor, M., Poole, M. et al. (2006) A naturally occurring epigenetic mutation in a gene encoding an SBP-box transcription factor inhibits tomato fruit ripening. Nature Genet. 38, 948–52.

Marí-Ordóñez, A., Marchais, A., Etcheverry, M., Martin, A., Colot, V. & Voinnet, O. (2013). Reconstructing de novo silencing of an active plant retrotransposon. Nature Genet. 45(9), 1029–39.

Matzke, M. A. & Mosher, R. A. (2014). RNA-directed DNA methylation: an epigenetic pathway of increasing complexity, Nat. Rev. Genet. 15, 394–408.

Maumus, F., Epert, A., Nogue, F. & Blanc, G. (2014). Plant genomes enclose footprints of past infections by giant virus relatives. Nature Commun. 5, 4268.

McKenna, A., et al. (2010) The Genome Analysis Toolkit: a MapReduce framework for analyzing next-generation DNA sequencing data. Genome research 20.9 1297–1303

Medrano, M., Herrera, C. M. & Bazaga P. (2014). Epigenetic variation predicts regional and local intraspecific functional diversity in a perennial herb. Mol. Ecol. 23, 4926–4938.

Mirouze, M., Reinders, J., Bucher, E., Nishimura, T., Schneeberger, K., Ossowski, S., Cao, J., Weigel, D., Paszkowski, J. & Mathieu, O. (2009). Selective epigenetic control of retrotransposition in *Arabidopsis*. Nature 461, 427–430.

Miura, A., Yonebayashi, S., Watanabe, K., Toyama, T., Shimada, H. & Kakutani, T. (2001). Mobilization of transposons by a mutation abolishing full DNA methylation in *Arabidopsis*. Nature 411(6834), 212–4.

Molaro, A., Hodges, E., Fang, F., Song, Q., McCombie, W. R., Hannon, G. J. & Smith, A. D. (2011). Sperm methylation profiles reveal features of epigenetic inheritance and evolution in primates. Cell 146(6), 1029–41.

Narum, S. R., Buerkle, C. A., Davey, J. W., Miller, M. R. & Hohenlohe, P. A. (2013). Genotyping-by-sequencing in ecological and conservation genomics. Mol. Ecol. 22, 2841–2847.

Niederhuth, C.E.*, Bewick, A.J.*, Ji, L., Alabday, M., Kim, K.D., Li, Q., Rohr, N.A., Rambani, A., Burke, J.M., Udall, J.A., Egesi, C., Schmutz, J., Grimwood, J., Jackson, S.A., Springer, N.M., Schmitz, R.J. (2016). Widespread natural variation of DNA methylation within angiosperms. Genome Biol. 17, 194.

Noy-Malka, C., Yaari, R., Itzhaki, R., Mosquna, A., Auerbach Gershovitz, N., Katz, A. & Ohad, N. (2014). A single CMT methyltransferase homolog is involved in CHG DNA methylation and development of *Physcomitrella patens*. Pl. Mol. Biol. 84, 719–735.

O’Malley, R.C. & Ecker, J.R. (2012). Epiallelic variation in *Arabidopsis thaliana*. Cold Spring Harb. Symp. Quant. Biol. 77, 135–145.

Ossowski, S., Schneeberger, K., Lucas-Lledo, J. I., Warthmann, N., Clark, R. M., Shaw, R. G., Weigel, D. & Lynch, M. (2010). The rate and molecular spectrum of spontaneous mutations in *Arabidopsis thaliana*. Science 327, 92–94.

Park P. (2009). ChIP-seq: advantages and challenges of a maturing technology. Nat. Rev. Genet. 10, 669–80.

Paszkowski, J. & Grossniklaus, U. (2011). Selected aspects of transgenerational epigenetic inheritance and resetting in plants. Curr. Opin. Pl. Biol. 14, 195–203.

Pavey, S. A., Bernatchez, L., Aubin-Horth, N. & Landry, C. R. (2012). What is needed for next-generation ecological and evolutionary genomics? Trends Ecol. & Evol. 27.12, 673–678.

Pecinka, A. & Mittelsten Scheid, O. (2012). Stress-induced chromatin changes: a critical view on their heritability. Plant Cell Physiol. 53, 801–808.

Platt, A., Gugger, P. & Sork, V.L. (2015). Genome-wide signature of local adaptation linked to variable CpG methylation in oak populations. Mol. Ecol. 24, 3823–3830.

Ragunathan, K., Jih, G. & Moazed, D. (2014). Epigenetic inheritance uncoupled from sequence-specific recruitment. Science 348, doi: 10.1126/science.1258699.

Rasmann, S., De Vos, M., Casteel, C. L., Tian, D., Halitschke, R., Sun, J. Y., Agrawal, A. A., Felton, G. W. & Jander, G. (2012). Herbivory in the previous generation primes plants for enhanced insect resistance. Plant Physiol. 158, 854–863.

Reinders, J., Wulff, B. B., Mirouze, M., Marí-Ordóñez, A., Dapp, M., Rozhon, W., Bucher, E., Theiler, G. & Paszkowski, J. (2009). Compromised stability of DNA methylation and transposon immobilization in mosaic *Arabidopsis* epigenomes. Genes Dev. 23, 939–950.

Rendina González, A. P., Chrtek, J., Dobrev, P. I. Dumalasová, V., Fehrer, J., Mráz, P. & Latzel, V. (2016). Stress-induced memory alters growth of clonal off spring of white clover (*Trifolium repens)*, Am. J. Bot. 103 (9), 1567 – 1574.

Rensing, S. A., Lang, D., Zimmer, A. D., Terry, A., Salamov, A., Shapiro, H., Nishiyama, T., Perroud, P.-F., Lindquist, E. A., Kamisugi, Y., Tanahashi, T., Sakakibara, K., Fujita, T., Oishi, K., Shin-I, T., Kuroki, Y., Toyoda, A., Suzuki, Y., Hashimoto, S.-i., Yamaguchi, K., Sugano, S., Kohara, Y., Fujiyama, A., Anterola, A., Aoki, S., Ashton, N., Barbazuk, W. B., Barker, E., Bennetzen, J. L., Blankenship, R., Cho, S. H., Dutcher, S. K., Estelle, M., Fawcett, J. A., Gundlach, H., Hanada, K., Heyl, A., Hicks, K. A., Hughes, J., Lohr, M., Mayer, K., Melkozernov, A., Murata, T., Nelson, D. R., Pils, B., Prigge, M., Reiss, B., Renner, T., Rombauts, S., Rushton, P. J., Sanderfoot, A., Schween, G., Shiu, S.-H., Stueber, K., Theodoulou, F. L., Tu, H., Van de Peer, Y., Verrier, P. J., Waters, E., Wood, A., Yang, L., Cove, D., Cuming, A. C., Hasebe, M., Lucas, S., Mishler, B. D., Reski, R., Grigoriev, I. V., Quatrano, R.S. & Boore, J. L. (2008). The *Physcomitrella* genome reveals evolutionary insights into the conquest of land by plants. Science 319, 64–69.

Reyna-Lopez, G. E., Simpson, J. & Ruiz-Herrera, J. (1997). Differences in DNA methylation patterns are detectable during the dimorphic transition of fungi by amplification of restriction polymorphims. Mol. Genet. & Genomics 253, 703–710.

Richards, C. L., Bossdorf, O. & Verhoeven, K. J. F. (2010). Understanding natural epigenetic variation. New Phytol. 187, 562–564.

Richards, C. L., Schrey, A. W. & Pigliucci, M. (2012). Invasion of diverse habitats by few Japanese knotweed genotypes is correlated with high epigenetic differentiation. Ecol. Lett. 15, 1016–1025.

Richards, E. J. (2006). Inherited epigenetic variation—revisiting soft inheritance. Nat. Rev. Genet. 7, 395–401.

Robertson, M. H. & Richards, C.L. (2015). Non-genetic inheritance in evolutionary theory - the importance of plant studies. Non-Gen. Inherit. 2, 3–11.

Robertson, M. H., Schrey, A. W., Shayter, A., Moss, C. J. & Richards, C. L. In press. Genetic and epigenetic variation in *Spartina alterniflora* following the *Deepwater Horizon* oil spill. Special issue in Evolutionary Applications: Evolutionary Toxicology.

Sani, E., Herzyk, P., Perrella, G., Colot, V. & Amtmann, A. (2013). Hyperosmotic priming of *Arabidopsis* seedlings establish a long-term somatic memory accompanied by specific changes of the epigenome. Genome Biol. 14, R59.

Schield, D. R., Walsh, M. R., Card, D. C., Andrew, A. L., Adams, R. H. & Castoe, T. A. (2015). EpiRADseq: scalable analysis of genomewide patterns of methylation using next-generation sequencing. Meth. in Ecol. & Evol. doi: 10.1111/2041-210X.12435.

Schmitz, R. J., Schultz, M. D., Urich, M. A., Nery, J. R., Pelizzola, M., Libiger, O., Alix A., McCosh R. B., Chen H. & Schork N. J. (2013). Patterns of population epigenomic diversity. Nature 495, 193–198 (2013).

Schrey, A.W., Alvarez, M., Foust, C.M., Kilvitis, H.J., Lee, J.D., Liebl, A.L., Martin, L.B., Richards, C.L. & Robertson M.H. (2013). Ecological Epigenetics: Beyond MS-AFLP. Integr. Comp. Biol. 53, 340–350.

Schulz, B., Eckstein, R. L. & Durka, W. (2014). Epigenetic variation reflects dynamic habitat conditions in a rare floodplain herb. Mol. Ecol. 23, 3523–3537.

Secco, D., Wang, C., Shou, H., Schultz, M. D., Chiarenza, S., Nussaume, L., Ecker, J. R., Whelan, J. & Lister, R. (2015). Stress induced gene expression drives transient DNA methylation changes at adjacent repetitive elements. eLife 4.

Seymour, D. K., Koenig, D., Hagmann, J., Becker, C. & Weigel, D. (2014). Evolution of DNA methylation patterns in the Brassicaceae is driven by differences in genome organization. PLoS Genet. 10, e1004785.

Slatkin, M. (2009). Epigenetic inheritance and the missing heritability problem. Genetics 182(3), 845–850.

Slaughter, A., Daniel, X., Flors, V., Luna, E., Hohn, B. & Mauch-Mani, B. (2012). Descendants of primed *Arabidopsis* plants exhibit resistance to biotic stress. Plant Physiol. 158, 835–843.

Song, Q., Decato, B., Hong, E. E., Zhou, M., Fang, F., Qu, J., Garvin, T., Kessler, M., Zhou, J. & Smith, A. D. (2013). A reference methylome database and analysis pipeline to facilitate integrative and comparative epigenomics. PLoS One 8(12), e81148.

Springer N. M., Lisch, D. & Li, Q. (2016). Creating order from chaos: Epigenome dynamics in plants with complex genomes. Plant Cell 28, 314–325.

Takuno S., Ran, J. H. & Gaut, B.S. (2016). Evolutionary patterns of genic DNA methylation vary across land plants. Nature Plants 2, 15222.

Taudt, A., Colomé-Tatché, M. & Johannes, F. (2016). Genetic sources of population epigenomic variation. Nat. Rev. Genet. 17(6), 319–32.

Teixeira, F. K., Heredia, F., Sarazin, A., Roudier, F., Boccara, M., Ciaudo, C., Cruaud, C., Poulain, J., Berdasco, M., Fraga, M.F., Voinnet, O., Wincker, P., Esteller, M. & Colot, V. (2009). A role for RNAi in the selective correction of DNA methylation defects. Science 323(5921), 1600–4.

Trucchi, E., Mazzarella, A. B., Gilfillan, G. D., Romero, M. T. & Paun, O. (2016). BsRADseq screening DNA methylation in natural populations of non-model species. Mol. Ecol. 25, 1697–1713.

Vergeer, P. & Ouborg, N. J. (2012). Evidence for an epigenetic rolein inbreeding depression. Biol. Lett. 8, 798–801.

Verhoeven, K. J. F. & Preite, V. (2014). Epigenetic variation in asexually reproducing organisms. Evol., 68, 644–655.

Verhoeven, K. J. F., Jansen, J. J., van Dijk, P. J. & Biere, A. (2010). Stress-induced DNA methylation changes and their heritability in asexual dandelions. New Phyt. 185(4), 1108–1118.

Wang, J., Fan, C. (2014). A neutrality test for detecting selection on DNA methylation using single methylation polymorphism frequency spectrum. Genome Biol Evol. 7(1), 154–71. DOI: 10.1093/gbe/evu271

Wibowo, A.T., Becker, C., Marconi, G., Durr, J., Price, J., Hagmann, J., Papareddy, R., Kageyama, J., Becker, J., Weigel, D. & Gutierrez-Marcos, J. (2016). Hyperosmotic stress memory in *Arabidopsis* is mediated by distinct epigenetically labile sites in the genome and is restricted in the male germline by DNA glycosylase activity. eLife 5:e13546.

Widiez, T., Symeonidi, A., Luo, C., Lam, E., Lawton, M. & Rensing, S. A. (2014). The chromatin landscape of the moss *Physcomitrella patens* and its dynamics during development and drought stress. The Plant Journal: for cell and molecular biology 79, 67–81.

Willing, E.-M., Rawat, V., Mandáková, T., Maumus, F., James, G. V., Nordström, K. J. V., Becker, C., Warthmann, N., Chica, C., Szarzynska, B., Zytnicki, M., Albani, M. C., Kiefer, C., Bergonzi, S., Castaings, L., Mateos, J. L., Berns, M. C., Bujdoso, N., Piofczyk, T., de Lorenzo, L., Barrero-Sicilia, C., Mateos, I., Piednoël, M., Hagmann, J., Chen-Min-Tao R., Iglesias-Fernández, R., Schuster S. C., Alonso-Blanco, C., Roudier, F., Carbonero, P., Paz-Ares, J., Davis, S. J., Pecinka, A., Quesneville, H., Colot, V., Lysak M. A., Weigel, D., Coupland, G. & Schneeberger, K. (2015). Genome expansion of *Arabis alpina* linked with retrotransposition and reduced symmetric DNA methylation. Nature Plants 1, Article number: 14023.

Wilschut, R. A., Oplaat, C., Snoek, L. B., Kirschner, J. & Verhoeven, K.J.F. (2016). Natural epigenetic variation contributes to heritable flowering divergence in a widespread asexual dandelion lineage. Mol. Ecol. 25, 1759–1768.

Xie, H. J., Li, A. H., Liu, A. D., Dai, W. M., He, J. Y., Lin, S., Duan, H., Liu, L. L., Chen, S. G., Song, X. L., Valverde, B. E. & Qiang, S. (2015). ICE1 demethylation drives the range expansion of a plant invader through cold tolerance divergence. Mol. Ecol. 24, 835–850.

Zhang, Y.Y., Fischer, M., Colot, V. & Bossdorf, O. (2013). Epigenetic variation creates potential for evolution of plant phenotypic plasticity. New Phytol. 197(1), 314–322.

